# Embryonic Durotaxis: A Mechanical Framework for Understanding Cesarean Scar Pregnancy

**DOI:** 10.1101/2025.04.24.650347

**Authors:** Jialing Cao, Huanyu You, Jianwen Li, Shiping Yue, Bingqi Song, Hangyu Li, Jiaqi Wei, Jine Zhang, Jiyu Wei, Zheng Guo, Dongshi Guan, Yan Zu, Zheng Xu, Yan Wang, Yubo Fan, Jing Du

## Abstract

Cesarean scar pregnancy (CSP) is an increasingly common complication associated with the rise in cesarean section rates, yet its underlying mechanisms remain poorly understood. In this study, we identified a stiffness gradient between healthy and scarred uterine tissues in both human cesarean samples and mouse uterine scar models. By employing gradient stiffness hydrogels, we demonstrated that mouse embryos and human trophoblast spheroids exhibit durotaxis, migrating preferentially toward stiffer areas. This migration occurs through three-dimensional complex behaviors—translation, swinging, and rolling—characterized by periodic patterns that align with cavity oscillations. Embryonic durotaxis is initiated from asymmetric adhesion forces and is driven by embryonic protrusive forces, generated from Marangoni-like tissue flows. The intrinsic cavity oscillations further amplify and sustain embryonic durotaxis. Finally, pharmacological inhibition of embryonic durotaxis significantly reduced CSP incidence in the mouse model. Our findings establish the concept of embryonic durotaxis, which may provide novel therapeutic avenues for CSP prevention.

## INTRODUCTION

The global increase in cesarean section (CS) rates has paralleled a rise in cesarean scar pregnancies (CSP), a serious and potentially life-threatening complication (*1, 2*). Clinical studies have shown that women with a history of CS face a heightened risk of embryo implantation within the cesarean scar (*3*), and this risk escalates with the number of prior CS procedures (*2*). When embryos implant in scarred tissue rather than normal endometrium (i.e., scar implantation), they can develop into CSP - a condition associated with severe outcomes, including uterine rupture, life-threatening hemorrhage, and major maternal morbidity as the pregnancy advances (*4, 5*). Although CSP is increasingly recognized in clinical settings for its grave implications, the fundamental mechanisms driving scar implantation over healthy tissue remain poorly understood (*6, 7*).

Embryo implantation occurs in three overlapping stages: apposition, adhesion, and invasion (*8, 9*). Of these, apposition—the stage during which the embryo rolls along the uterine lining—is critical in determining where implantation ultimately occurs (*8, 9*). Most current investigations into CSP pathogenesis focus on the severe progression of the disease (*10, 11*). Several factors, such as localized hypoxia, disrupted vascular remodeling, and altered immune responses, have been implicated at the scar site (*12*). However, a central unanswered question remains: why do embryos favor implantation in scarred tissue in the first place? This knowledge gap may hold the key to effective CSP prevention strategies.

Embryo movement in mammals has been attributed to uterine smooth muscle contractions that drive pre-implantation clustered migration (*13*). In humans, uterine contractions and intraluminal fluid flow also regulate embryo transport, and abnormal pressure gradients, such as those caused by hydrosalpinx, can displace embryos from implantation-competent regions (*14*). External mechanical constraints from maternal tissues further influence development by triggering anterior–posterior axis formation(*15*). More recently, embryos themselves have been shown to generate traction forces and remodel their surrounding matrix in a species-specific manner (*16*). Together, these studies highlight the impact of maternal mechanical cues and embryonic force generation on early development, yet whether embryos can perform directed, mechanosensitive migration remains unknown.

Recent research suggests that altered tissue biomechanics can actively drive disease by promoting pathological remodeling, such as fibrosis, impaired repair, and even tumor progression (*17–19*). Previous studies confirmed that cesarean scars are mechanically distinct, displaying significantly higher stiffness than the adjacent myometrium, with a reported strain ratio of 1.8 ± 0.7 (*20*). Yet, despite these insights, the potential contribution of uterine mechanical changes to CSP pathogenesis remains largely unexplored. Many cell types exhibit durotaxis, a mechanosensitive behavior in which cells migrate individually or collectively toward stiffer substrate regions (*21–27*). Our previous work demonstrated that human trophoblast cells migrate durotactically on substrates with stiffness gradients, suggesting that mechanical cues regulate trophoblast migration and may guide embryo implantation (*28*). Given that cesarean scar tissue exhibits markedly increased stiffness compared to surrounding healthy uterine tissue (*20, 29*), embryonic durotaxis may play a role in CSP development. However, whether mammalian embryos demonstrate durotaxis, and how this mechanotactic behavior affects CSP pathogenesis, remains unclear and requires further study.

In this study, we first characterized the mechanical properties of uterine scars in a murine CSP model and human cesarean samples, revealing a stiffness gradient that increases from healthy to scarred tissue. We then engineered polyacrylamide hydrogels with tunable stiffness gradients to replicate the mechanical environment of cesarean scars. Both peri-implantation stage mouse embryos and human trophoblast spheroids exhibited durotactic migration on these substrates. Combining experimental observations with mechanical modeling, we found that embryonic durotaxis initiates from asymmetric substrate adhesion and is primarily driven by polarized protrusive forces. This process involves diverse 3D modes of movement, including translation, swinging, and rolling, and features a distinct periodicity that synchronizes with blastocyst cavity oscillations. Finally, using our established uterine surgical scar mouse model, we demonstrated that pharmacological inhibition of embryonic protrusions significantly reduces CSP incidence. Our findings indicate that embryos are not only passively displaced by maternal forces but can also actively migrate in response to mechanical cues, exhibiting durotaxis, highlighting a promising therapeutic strategy for preventing this high-risk obstetric complication.

## RESULTS

### Mouse embryos and human trophoblast spheroids exhibit durotactic migration

To examine how cesarean sections influence embryo implantation, we established a murine model of uterine surgical scarring based on previously described protocols (*30, 31*) (Fig. 1A). Histological analysis confirmed that our model replicates key features of cesarean-induced myometrial disruption and subsequent scar formation (Supplementary Fig. S1A–C). Quantification of implantation sites revealed a significantly higher proportion of embryos implanted within scarred tissue compared to adjacent non-scarred regions (Fig. 1B and C). Importantly, embryos implanted in scarred areas exhibited poorer pregnancy outcomes, including increased rates of pregnancy loss and embryonic resorption in mice subjected to metratomy (Supplementary Fig. S1D and E). These results validate the utility of our murine model for investigating CSP pathogenesis and highlight the clinical significance of scar-associated implantation failure.

**Figure 1.**
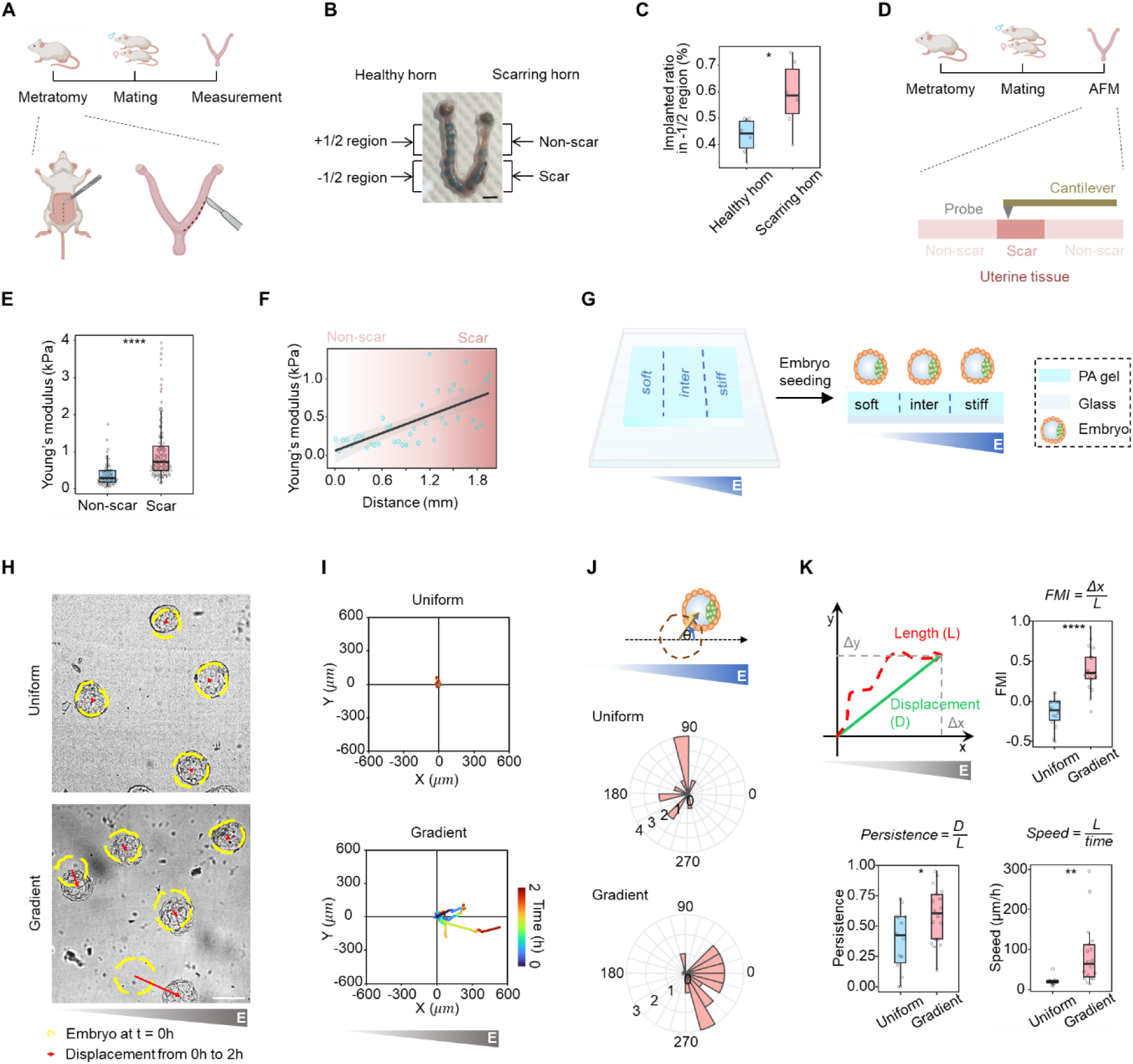
Stiffness gradient in the scarred murine uterine and durotaxis of embryos. (**A**) Schematic illustration of murine uterine scar model. (**B**) The image of implantation sites in murine uterine scar model. Scale bar: 5 mm. (**C**) The statistical analysis of implantation site percentage in - 1/2 region at healthy horn or scar horn of murine uterine scar model, n = 6. (**D**) Schematic illustration of AFM experiments in the scarred murine uterine. (**E**) The statistical analysis of Young’s modulus in scarred regions and non-scarred regions in the scarred murine uterine, n = 3. (**F**) The AFM analysis of Young’s modulus from non-scarred to scarred regions in the scarred murine uterine. (**G**) Schematic illustration of polyacrylamide hydrogel substrate system featuring a controlled stiffness gradient. (**H**) Representative time-lapse images of embryos on substrates with uniform stiffness or gradient stiffness. Scale bar: 200 μm. (**I**) Embryo migration plots on gradient substrates over 2 h. (**J**) Rose diagram of embryo migration direction on substrates with uniform stiffness or gradient stiffness, which displays the angular between migration and stiffness gradient and the frequency of each class. (**K**) The statistical analysis of durotaxis-related parameters, including Forward Migration Index (FMI), persistence, and velocity of embryo migration on substrates with uniform stiffness or gradient stiffness, n = 14. All box- and-whisker plots show the medians, maxima, minima, upper quartiles, and lower quartiles. * p < 0.05, ** p < 0.01, *** p < 0.001, **** p < 0.0001.

To explore the contribution of uterine mechanical properties to CSP, we used Atomic Force Microscopy (AFM) to assess tissue stiffness in both scarred and adjacent healthy regions at embryonic day 3.5 (E3.5; Fig. 1D). AFM measurements revealed that scarred tissue exhibited significantly higher stiffness than surrounding non-scarred regions (Fig. 1E). We also identified a clear stiffness gradient, with the Young’s modulus progressively increasing from non-scarred to scarred tissue (Fig. 1F). Similar stiffness gradient was also observed in human post-cesarean uterine samples from non-scarred to scarred tissue (Supplementary Fig. S1F). These results establish a direct correlation between tissue stiffness gradients and the preferential implantation of embryos in scarred regions.

To assess how embryos respond to the stiffness gradient characteristic of scarred uterine tissue, we cultured peri-implantation stage mouse embryos (E4.5) on collagen I-coated polyacrylamide hydrogels engineered with a defined stiffness gradient - hereafter referred to as gradient substrates - to simulate the mechanical environment of cesarean scars (*28*) (Fig. 1G). Time-lapse imaging revealed that embryos underwent robust durotactic migration, consistently moving from softer toward stiffer regions, albeit with varying velocities. In contrast, embryos placed on hydrogels with uniform stiffness - uniform substrates - exhibited no directional migration (Fig. 1H–J and Supplementary Video S1). Quantitative analysis of key durotaxis metrics, including Forward Migration Index (FMI), persistence, and velocity (*30, 32*) confirmed significantly greater durotactic activity on gradient substrates compared to uniform controls (Fig. 1K). When we further analyzed these parameters across distinct zones—soft, intermediate, and stiff—the strongest durotactic responses occurred in intermediate stiffness regions (Supplementary Fig. S2 and Supplementary Video S2). These results demonstrate that mouse embryos actively sense and respond to mechanical cues by migrating toward regions with higher stiffness.

In our previous work, we reported durotactic behavior in individual human trophoblast cells (JAR line) (*28*). Here, we extended our investigation to JAR multicellular spheroids, which also exhibited directed migration toward stiffer regions on gradient substrates (Supplementary Fig. S3A–F). However, comparative quantification of durotaxis parameters revealed that mouse embryos exhibited significantly stronger durotactic responses than JAR spheroids under identical gradient conditions (Supplementary Fig. S3G–I). Together, these findings support a mechanobiological basis for embryo guidance in the uterine environment and underscore the potential role of tissue stiffness gradients in driving embryonic migration during CSP pathogenesis.

### Spatial asymmetry in trophectoderm-substrate adhesion triggers embryonic durotaxis

Extensive research has established that cell-substrate interactions play a crucial role in driving durotaxis in both individual cells and cell collectives (*33–36*). In our study, we examined embryo-substrate adhesion by tracking trophectoderm cell membranes at the embryo-substrate interface using mTmG^+/-^ embryos. Time-lapse imaging revealed that upon initial contact, membrane signals concentrated predominantly on the stiffer side of the substrate, reflecting an asymmetric adhesion pattern aligned with the stiffness gradient (Fig. 2A–D and Supplementary Video S3). For clarity, all references to the “soft side” and “stiff side” of the embryo denote its orientation relative to the substrate stiffness gradient, not intrinsic mechanical differences within the embryo. As adhesion matured, membrane signal intensity increased significantly more on the stiff side than on the soft side, amplifying this asymmetry over time (Fig. 2E). We observed a similar asymmetric adhesion pattern in JAR spheroids cultured on gradient substrates, where Vinculin-GFP and Lifeact-mCherry signals displayed polarized spatial distributions at the interface (Supplementary Fig. S4A–D). Together, these results demonstrate that embryos establish distinct, spatially asymmetric trophectoderm-substrate adhesions in response to stiffness gradients, which likely underpin durotactic migration.

**Figure 2.**
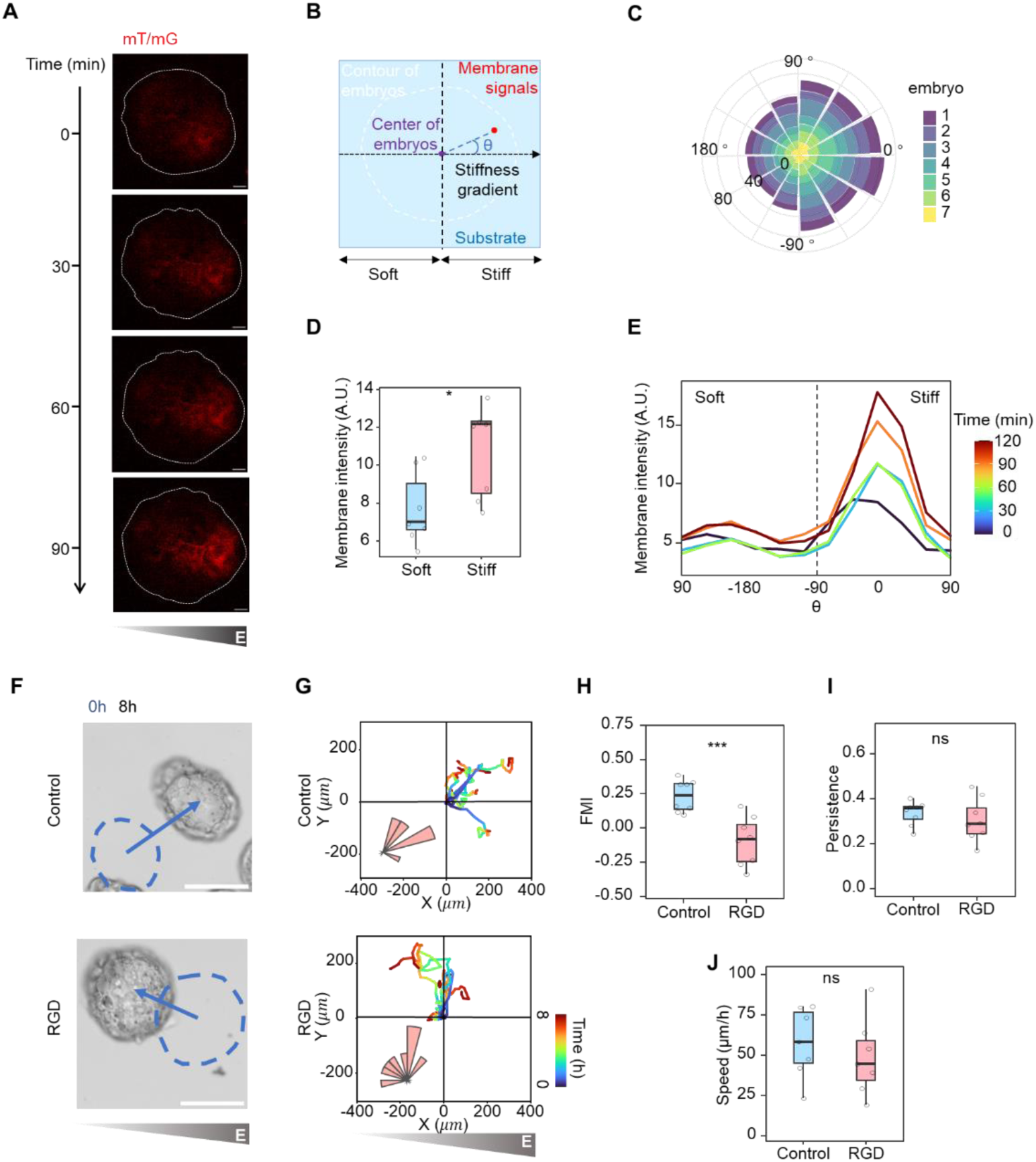
The establishment of an asymmetric trophectoderm-substrate adhesion pattern initiates embryonic durotaxis. (**A**) Representative time-lapse fluorescent images of trophectoderm cell membrane at the embryo-substrate interface in mTmG^+/-^ embryos. Scale bar: 20 μm. (**B**) Schematic illustration of angles (θ) between membrane signal related to embryo center and substrate stiffness gradient. (**C**) Rose diagram of membrane signals at the embryo-substrate interface in mTmG^+/-^embryos, which displays angles (θ) between membrane signal related to embryo center and substrate stiffness gradient. (**D**) The membrane intensity at the embryo-substrate interface in mTmG^+/-^ embryos at different angles (θ) during the adhesion of embryos to substrates. (**E**) The statistical analysis of membrane intensity at the embryo-substrate interface in soft side (θ: from 90⁰ to −90⁰) and stiff side (θ: from −90⁰ to 90⁰) of mTmG^+/-^ embryos, n = 7. (**F**) Representative time-lapse images of embryos on stiffness gradient substrates treated with control or RGD. Scale bar: 100 μm. (**G**) Embryo migration plots on stiffness gradient substrates over 8 h treated with control or RGD. Insets are rose diagram of embryo migration direction, which displays the angular between migration and stiffness gradient and the frequency of each class. (**H-J**) The statistical analysis of durotaxis-related parameters, including Forward Migration Index (FMI), persistence, and velocity of embryo migration on stiffness gradient substrates over 8 h treated with control or RGD, n = 8. All box-and-whisker plots show the medians, maxima, minima, upper quartiles, and lower quartiles. * p < 0.05, ** p < 0.01, *** p < 0.001, **** p < 0.0001.

To assess whether asymmetric trophectoderm-substrate adhesion is essential for embryonic durotaxis, we disrupted this interaction pharmacologically using RGD peptides(*37*), known inhibitors of integrin-mediated adhesion. RGD treatment markedly reduced the spatial asymmetry of focal adhesion (FA) distribution at the trophectoderm-substrate interface (Supplementary Fig. S4E, F). Quantitative analysis of durotaxis parameters - including Forward Migration Index, persistence, and velocity - revealed significantly impaired durotactic responses in RGD-treated embryos compared to DMSO-treated controls (Fig. 2F–J). Similarly, RGD-treated JAR spheroids exhibited reduced durotactic migration (Supplementary Fig. S4G–I). Given that adhesion is the first step in embryo-substrate interaction, these results confirm that spatially asymmetric trophectoderm adhesion is a critical prerequisite for initiating embryonic durotaxis.

### Polarized trophectoderm protrusions induced by Marangoni-like tissue flows drive embryonic durotaxis

Next, to investigate trophectoderm cell dynamics during embryonic migration, we compared embryo surface contours obtained via time-lapse imaging on gradient versus uniform substrates. Embryos on gradient substrates displayed significantly more dynamic and asymmetric protrusions than those on uniform substrates. These protrusions developed progressively and became more pronounced on the stiff side, indicating polarized surface activity aligned with the substrate stiffness gradient (Fig. 3A–F). To further characterize these polarized protrusions, we performed surface curvature analysis on mTmG^+/-^ embryos, which revealed a striking accumulation of cell blebs specifically localized to the stiff side of the embryonic surface (Fig. 3G). Quantitative analysis revealed a significant alignment between the orientation of dominant blebs and the direction of embryonic movement (Fig. 3H-J and Supplementary Video S4). Similar results were observed in EYFP-mem^+^ JAR spheroids (Supplementary Fig. S5). Together, these findings demonstrate that polarized protrusive activity closely correlates with the directionality of embryonic durotaxis, reinforcing the role of mechanical sensing in guiding embryo migration.

**Figure 3.**
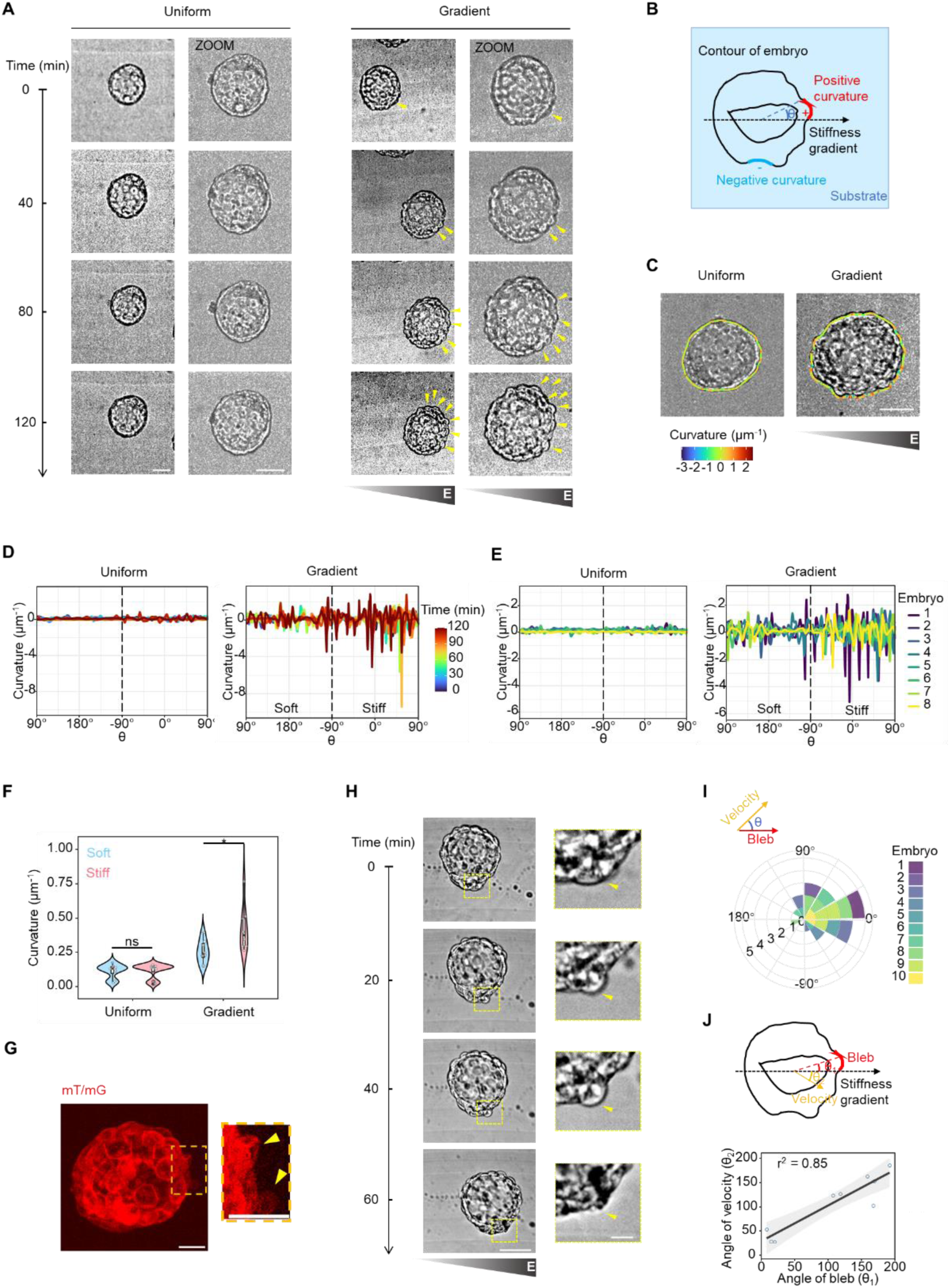
Polarized surface fluctuations during embryonic durotaxis on stiffness gradient substrates. (**A**) Representative time-lapse images of embryo surface morphology over time on substrates with uniform stiffness or gradient stiffness. The yellow arrows indicate trophectoderm protrusions. Scale bar: 50 μm. (**B**) Schematic illustration of angles (θ) between surface curvature related to embryo center and substrate stiffness gradient. (**C**) The analysis of surface curvature of embryos on substrates with uniform stiffness or gradient stiffness. Scale bar: 50 μm. (**D**) The surface curvature of embryos at different angles (θ) over time on substrates with uniform stiffness or gradient stiffness. (**E**) The surface curvature of embryos at different angles (θ) on substrates with uniform stiffness or gradient stiffness. (**F**) The statistical analysis of surface curvature in soft side (θ: from 90⁰ to −90⁰) and stiff side (θ: from −90⁰ to 90⁰) of embryos on substrates with uniform stiffness or gradient stiffness, n = 8. (**G**) Representative fluorescent images of surface morphology highlighting an active cell bleb. Scale bar: 20 μm. (**H**) Representative time-lapse images of embryo surface morphology over time on substrates with uniform stiffness or gradient stiffness. The yellow arrows indicate trophectoderm protrusions. Scale bar: 50 μm. (**I**) Rose diagram displays the angular between the orientation of dominant blebs and the direction of embryonic movement during embryo durotaxis on stiffness gradient substrates. (**J**) The correlation analysis of the orientation of dominant blebs and the direction of embryonic movement, n = 8. All box-and-whisker plots show the medians, maxima, minima, upper quartiles, and lower quartiles. * p < 0.05, ** p < 0.01, *** p < 0.001, **** p < 0.0001.

To better understand how polarized protrusions form during durotactic migration in embryos, we labeled cell nuclei with SPY555 and performed time-lapse imaging. This analysis revealed a markedly higher density of trophectoderm cells on the stiff side of embryos compared to the soft side, indicating the emergence of polarized cellular crowding during migration (Fig. 4A–C). Morphological analysis further revealed that the nuclei of trophectoderm cells on the stiff substrate appeared more flattened than those on the soft side (Fig. 4A and D), indicating that substrate stiffness gradients induce directional nuclear deformation.

**Figure 4.**
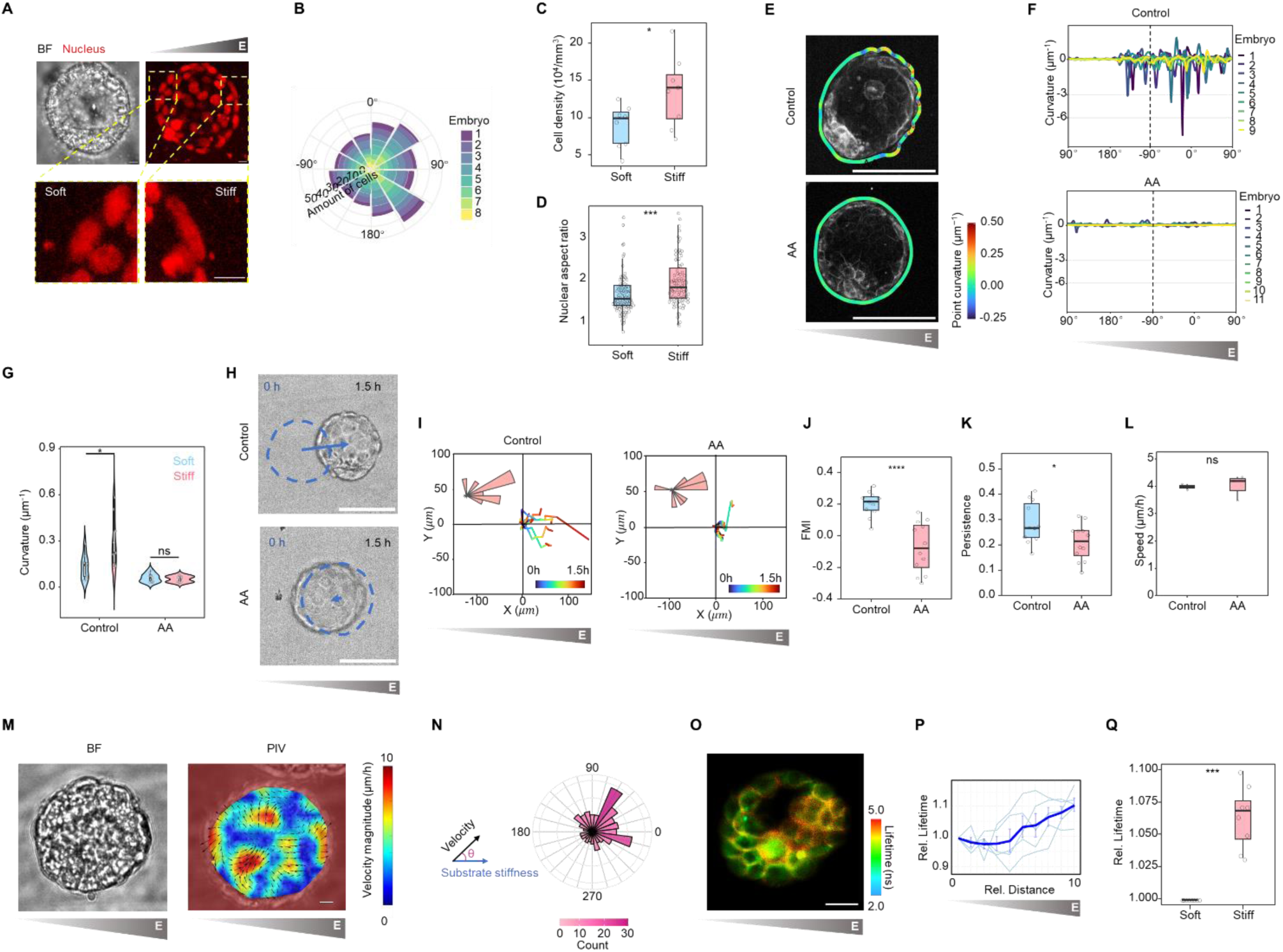
Marangoni-like tissue flow-induced cell blebbing is essential for embryonic durotaxis. (**A**) Representative fluorescent images of cell nuclei labeled by SPY555 during embryonic durotaxis on stiffness gradient substrates. Scale bar: 15 μm. (**B**) The cell density at different angles (θ) on substrates with gradient stiffness. (**C**) The statistical analysis of cell number in soft side (θ: from 90⁰ to −90⁰) and stiff side (θ: from −90⁰ to 90⁰) of embryos on stiffness gradient substrates, n = 8. (**D**) The statistical analysis of nuclear aspect ratio in soft side (θ: from 90⁰ to −90⁰) and stiff side (θ: from −90⁰ to 90⁰) of embryos on stiffness gradient substrates, n = 8. (**E**) The analysis of embryo surface curvature on stiffness gradient substrates treated with control or AACOCF3. Scale bar: 100 μm. (**F**) The surface curvature of embryos at different angles (θ) on stiffness gradient substrates treated with control or AACOCF3. (**G**) The statistical analysis of surface curvature in soft side (θ: from 90⁰ to −90⁰) and stiff side (θ: from −90⁰ to 90⁰) of embryos on stiffness gradient substrates treated with control or AACOCF3, n = 9 (control), 11 (AA). (**H**) Representative time-lapse images of embryos on stiffness gradient substrates treated with control or AACOCF3. Scale bar: 100 μm. (**I**) Embryo migration plots on stiffness gradient substrates over 1.5 h treated with control or AACOCF3. Insets are rose diagram of embryo migration direction, which displays the angular between migration and stiffness gradient and the frequency of each class. (**J-L**) The statistical analysis of durotaxis-related parameters, including Forward Migration Index (FMI), persistence, and velocity of embryo migration on stiffness gradient substrates over 1.5 h treated with control or AACOCF3, n = 12. (**M**) Representative velocity field of embryos during durotaxis on stiffness gradient substrates. Scale bar: 50 μm. (**N**) Rose diagram displays the angular between velocity field and substrate stiffness gradient. (**O**) Representative FLIM images of plasma membrane tension probed by Flipper-TR on the embryo surface close to the embryo-substrate interface. Scale bar, 30 μm. (P)The relationship between relative fluorescence lifetime of Flipper-TR and relatvie distance from soft side of the embryo, n = 5. (**Q**) Relative fluorescence lifetime of Flipper-TR after selecting the plasma membrane as the region of interest (ROI) on soft and stiff sides of embryo, n = 8. All box-and-whisker plots show the medians, maxima, minima, upper quartiles, and lower quartiles. * p < 0.05, ** p < 0.01, *** p < 0.001, **** p < 0.0001.

Previous studies have identified nuclear deformation as a key driver of cell blebbing through cytosolic phospholipase A2 (cPLA2) signaling pathways (*38*). To examine this mechanism in our model, we treated embryos with AACOCF3, a potent and selective cPLA2 inhibitor (*38, 39*). This treatment significantly reduced the number of trophectoderm protrusions and disrupted their spatial polarity within embryos (Fig. 4E–G). Quantitative analysis of parameters associated with durotaxis revealed that AACOCF3 markedly impaired embryonic durotaxis (Fig. 4H–L and Supplementary Video S5). We confirmed these findings in parallel experiments using EGFP-NLS⁺ and EYFP-mem⁺ JAR spheroids, which produced consistent results (Supplementary Fig. S5). Together, these data demonstrate that polarized trophectoderm protrusions, driven by nuclear deformation in response to substrate stiffness gradients, play a critical role in guiding embryonic durotaxis.

To uncover the biophysical mechanism behind polarized embryonic protrusions on stiffness gradients, we applied Particle Image Velocimetry (PIV) to analyze tissue flows. This revealed directional internal flows toward the stiff side of embryos, which strongly correlated with protrusion formation (Fig. 4M and N). These flows corresponded with the increased cell density observed on the stiff side (Fig. 4A–C). Recent work has highlighted the role of surface tension in driving tissue flow (*40, 41*). To investigate this, we examined surface tension differences between the stiff and soft sides of embryos on gradient substrates by FLIM imaging. We found a gradient increase in fluorescence lifetime of Flipper-TR on the PM align with the substrate stiffness gradient (Fig. 4O–Q). These results support a Marangoni-like flow mechanism (*40, 41*), where the trophectoderm cells first establish polarized adhesion patterns aligned with the gradient and produces surface tension gradient which then generate internal flows: elevated tension on the stiff side drives cells inward, increasing local cell density and enhancing protrusive activity.

### The 3D interplay between protrusive forces and substrate adhesion orchestrates multi-modal embryonic durotaxis

To elucidate the mechanisms driving durotactic migration in embryos, we employed live-cell fluorescent dye labeling to visualize nuclei and track cell movements in real-time. Interestingly, unlike single cells migrating on 2D substrates - which typically undergo only translational motion without rotation - mouse embryos exhibited multiple modes of durotactic movement: (1) translation, (2) swing, characterized by in-plane rotation around an axis perpendicular to the substrate, and (3) rolling, defined as out-of-plane rotation around an axis parallel to the substrate (Fig. 5A and Supplementary Video S6– S8). Quantitative analyses showed that swing was the dominant migration mode across all substrate stiffness zones (soft, intermediate, and stiff). The distribution of these migration modes varied significantly depending on local stiffness: swing predominated on substrates of intermediate stiffness, while translation occurred more frequently on stiff regions. In contrast, rolling was more common on soft areas (Fig. 5B), demonstrating a precise mechanosensitive regulation of migratory patterning.

**Figure 5.**
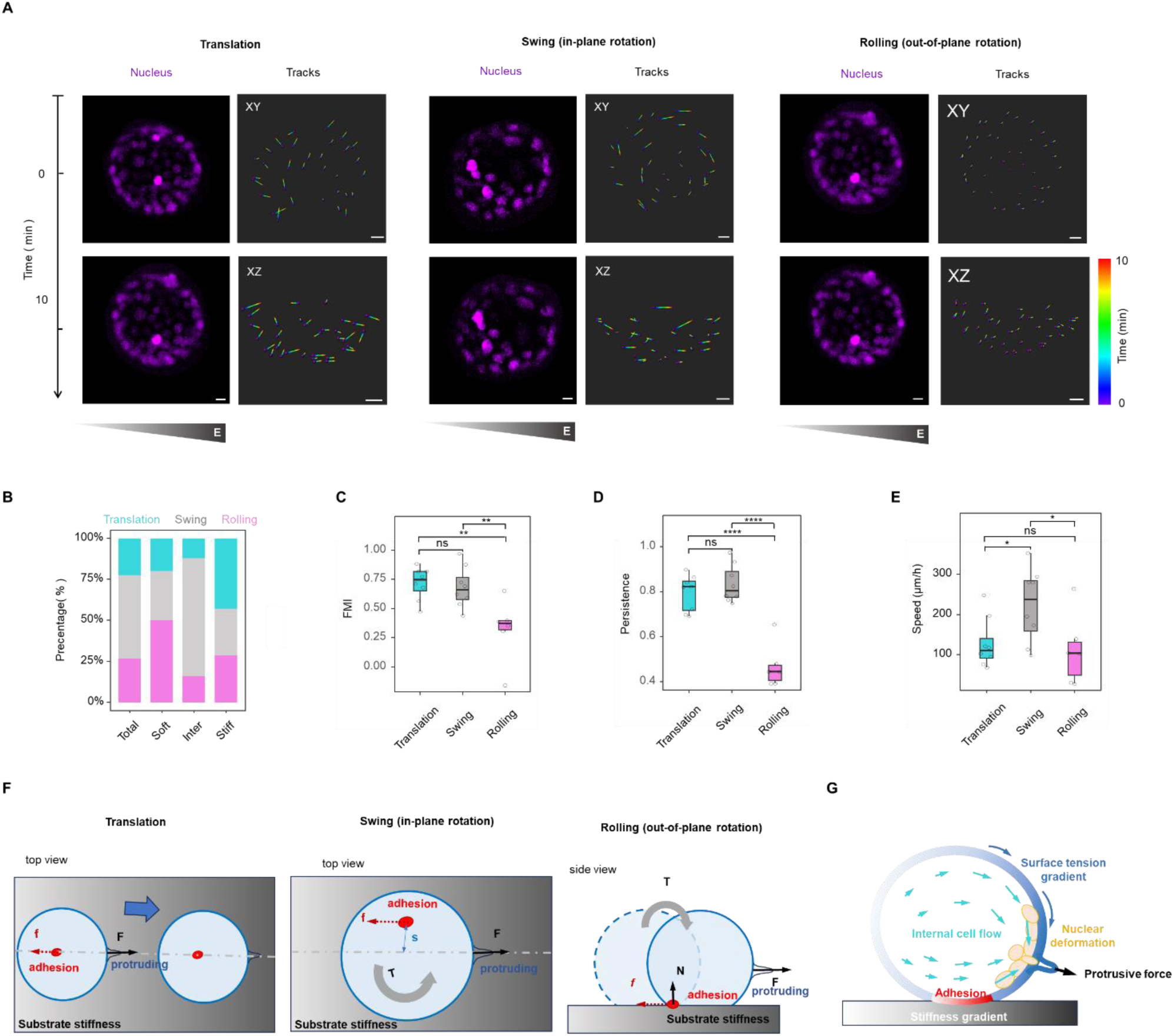
Multiple migration modes of embryonic durotaxis driven by 3D interaction between protrusive forces and substrate adhesion. (**A**) Representative fluorescent images and spatiotemporal trajectory of cell nuclei labeled by SPY555 during embryonic durotaxis on stiffness gradient substrates. Scale bar: 50 μm. (**B**) The distribution of durotactic modes across different stiffness zones (total: n = 49, soft: n=10, intermediate: n = 25, stiff: n = 14), (**C-E**) The statistical analysis of durotaxis-related parameters, including Forward Migration Index (FMI), persistence, and velocity of three embryo migration modes on stiffness gradient substrates, n = 8. (**F**) Schematic diagram of force analysis on embryos under three migration modes. (**G**) Schematic diagram of protrusive force-driven embryo durotaxis under Marangoni-like tissue flows. All box-and-whisker plots show the medians, maxima, minima, upper quartiles, and lower quartiles. * p < 0.05, ** p < 0.01, *** p < 0.001, **** p < 0.0001.

To assess the efficiency of each migration mode, we measured FMI, persistence, and velocity. Both translation and swing modes exhibited greater migratory efficiency than rolling, as evidenced by higher FMI, persistence, and speed (Fig. 5C–E). Among all modes, swing resulted in the highest migration velocity (Fig. 5E). Together, these results indicate that mouse embryos employ a dynamic repertoire of migration strategies—coordinating translation, swing, and rolling—to navigate stiffness gradients efficiently during durotaxis.

To explore how polarized protrusive forces drive distinct migration modes, we constructed a mechanical model based on our experimental findings, considering both active protrusive force and substrate adhesion force. Because cell blebbing and embryonic migration were highly dynamic, we treated them as nonequilibrium processes. In the translational mode, trophectoderm protrusions form predominantly at the embryo’s basal surface, generating a protrusive force *F* that opposes the substrate drag force *f* along the same axis. When *F* and *f* become misaligned, their separation generates a torque T, which induces either swing (in-plane rotation) or rolling (out-of-plane rotation) (Fig. 5F). This model mechanistically accounts for the greater efficiency observed in the swing mode (Fig. 5C–E). Translation requires that *F* overcomes the maximum drag force *fₘ*(*F* ≥ fₘ), where *fₘ* primarily results from embryo-substrate adhesion. In contrast, rotational modes—swing and rolling—can still occur when *F* is less than *fₘ*, as they are governed by torque dynamics rather than linear force balance.

According to our model, torque is described by *T = Fs = 8πμR³ω*, where *s* is the separation between *F* and *f*, μ is fluid viscosity, *R* is embryo radius, and ω is angular velocity. Because rolling also requires detachment from the substrate in addition to torque generation, swing emerges as the more energy-efficient and more frequent mode - aligning with our experimental observations (Fig. 5B–E).

Our data further emphasize that substrate stiffness plays a key role in determining the preferred mode of migration (Fig. 5B). On stiffer substrates, strong adhesion aligns F and f, promoting translation. In contrast, intermediate and soft substrates disrupt this alignment, favoring rotational motion. On soft substrates, weak adhesion fails to resist out-of-plane torque, thereby facilitating rolling. Collectively, these results demonstrate that substrate stiffness regulates the balance between protrusive and adhesive forces, thereby dictating which durotactic migration mode embryos predominantly adopt. We proposed a mechanism that drives embryonic durotaxis: polarized trophectoderm adhesion on substrate generates a surface tension gradient, which induces Marangoni-like tissue flows directed toward the stiff side, leading to polarized trophectoderm protrusions. The 3D interaction between protrusive forces and substrate adhesion drives different models of embryonic durotaxis (Fig. 5G). This model directly connects substrate stiffness gradients to internal cell movement and protrusions, offering a unified framework for understanding embryonic durotaxis.

### Intrinsic cavity oscillations amplify and sustain embryonic durotaxis

As previously noted, mouse embryos exhibited substantially stronger durotaxis than JAR spheroids (Supplementary Fig. S3G–I). By comparing their migration trajectories, we found that while JAR spheroids showed only brief and limited durotactic responses, mouse embryos engaged in periodic and sustained durotactic migration (Fig. 6A and Supplementary Video S9). Prior studies have described the cyclic expansion and contraction of the mammalian blastocyst cavity, a phenomenon known as cavity oscillations (*42, 43*). Our quantitative analysis revealed a tight temporal correlation between these oscillatory dynamics and embryonic migration cycles (Fig. 6B and C). Within each oscillation cycle, embryos underwent continuous and directed migration during cavity expansion phases. In contrast, durotactic movement was disrupted during contraction phases (Fig. 6B and D). These findings highlight a distinctive migratory behavior in mouse embryos, where intrinsic cavity oscillations modulate embryonic migration in response to external mechanical gradients.

**Figure 6.**
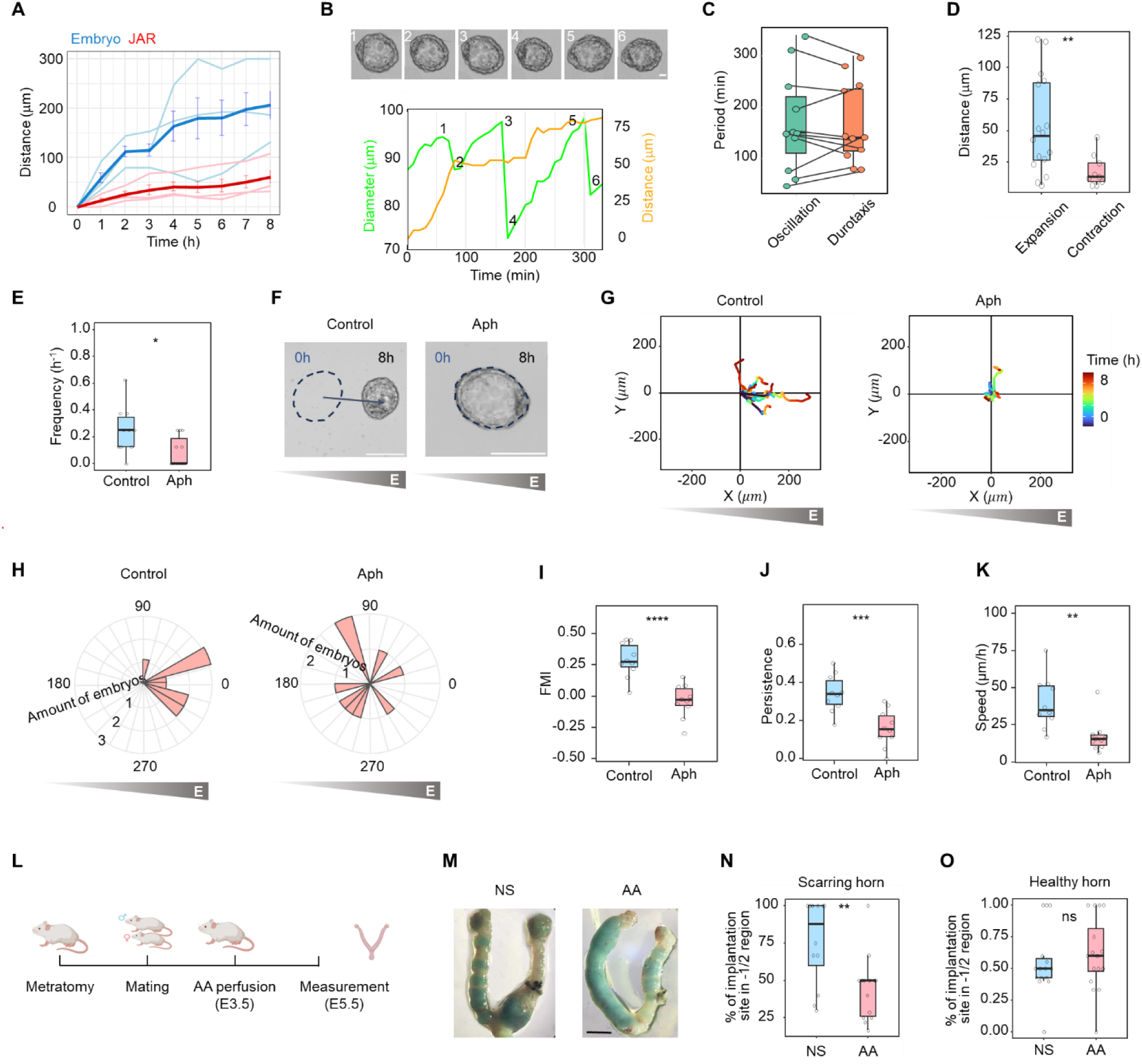
Durotaxis of mouse embryos exhibits periodicity and contributes to CSP pathogenesis. (**A**) The relationship between the distance to the endpoint and migration time during durotaxis of a JAR spheroid or a mouse embryo. (**B**) The relationships between the distance to the endpoint and migration time or the embryo diameter and migration time during durotaxis of a mouse embryo. (**C**) The statistical analysis of period duration in embryo oscillation or durotaxis, n = 11. (**D**) The statistical analysis of migration length in expansion phase or contraction phase in a single oscillation period, n = 16. (**E**) Oscillation frequency of embryos treated with control or aphidicolin, n = 10 (control), 11 (Aph). (**F**) Representative time-lapse images of embryos on stiffness gradient substrates treated with control or aphidicolin. Scale bar: 50 μm. (**G**) Migration plots on gradient substrates over 8 h in embryos treated with control or aphidicolin. (**H**) Rose diagram of migration direction on substrates with gradient stiffness in embryos treated with control or aphidicolin, which displays the angular between migration and stiffness gradient and the frequency of each class. (**I-K**) The statistical analysis of durotaxis-related parameters, including Forward Migration Index (FMI), persistence, and velocity of embryo migration on stiffness gradient substrates over 8 h treated with control or AACOCF3, n = 10. (**L**) Schematic illustration of AACOCF3 perfusion treatment in pregnant mice uterine scar model. (**M**) The image of implantation sites in murine uterine scar model treated with control (normal Saline, NS) or AACOCF3. Scale bar: 5 mm. (**N**) The statistical analysis of implantation site percentage in −1/2 region at scar horn of murine uterine scar model treated with control (normal Saline, NS) or AACOCF3, n = 12. (**O**) The statistical analysis of implantation site percentage at healthy horn of murine uterine scar model treated with control (normal Saline, NS) or AACOCF3, n = 16. All box-and-whisker plots show the medians, maxima, minima, upper quartiles, and lower quartiles. * p < 0.05, ** p < 0.01, *** p < 0.001, **** p < 0.0001.

To test the role of cavity oscillations in durotactic migration, we treated embryos with aphidicolin, which inhibits trophectoderm cell division, a process required to sustain cavity oscillations (*42*). Aphidicolin treatment significantly reduced oscillation frequency compared to vehicle-treated controls (Fig. 6E) and led to a marked decline in durotactic migration (Fig. 6F–H). Quantitative analysis revealed that all key metrics of durotaxis—FMI, persistence, and velocity—were significantly impaired following treatment (Fig. 6I–K). Together, these findings demonstrate that cavity oscillations amplify and sustain protrusive force–driven durotaxis by introducing rhythmic, periodic responses to stiffness gradients, thereby enabling embryos to achieve longer-range directed migration compared to JAR spheroids.

### Inhibiting embryonic durotaxis prevents ectopic implantation in cesarean scars

To directly examine the pathophysiological relevance of embryonic durotaxis in CSP, we applied AACOCF3 perfusion in our established murine CSP model featuring surgically induced uterine scars (Fig. 6L). Pharmacological inhibition of cell blebbing - and thus durotaxis - led to a marked reduction in implantation frequency within scarred regions, while implantation in adjacent non-scarred areas remained unaffected. Notably, AACOCF3 treatment did not alter implantation site distribution within the healthy uterine horn (Fig. 6M–O). These findings underscore the influence of local uterine mechanics in guiding embryo implantation, suggesting that targeting durotactic migration may offer a preventive strategy for CSP.

## DISCUSSION

Cesarean scar pregnancy is a growing obstetric concern with limited preventive strategies, mainly due to an incomplete understanding of its underlying pathogenesis - especially the factors that drive aberrant implantation within scarred uterine tissue (*44–47*). To address this gap, we first identified a distinct mechanical stiffness gradient spanning from non-scarred to scarred zones in both human post-cesarean uterine tissues and mouse uterine scar models. Using polyacrylamide (PA) hydrogels engineered with tunable stiffness, we replicated these mechanical gradients and systematically investigated their impact on embryo migration behavior.

Our study demonstrates that both mouse embryos and human trophoblast spheroids undergo directional migration – durotaxis - along defined substrate stiffness gradients. In this process, trophectoderm cells at the embryo–substrate interface first detect the stiffness gradient and establish polarized adhesion patterns aligned with the gradient. Further mechanistic analyses revealed that internal tissue flows directed toward the stiff side drive the formation of polarized trophectoderm protrusions. These protrusions, in turn, generate the forces necessary for durotaxis through three distinct kinematic modes: translation, in-plane swing, and out-of-plane rolling. We also discovered that the periodic expansion and contraction of the blastocyst cavity reinforce and prolong this migration behavior. Specifically, cavity oscillations synchronize with durotactic phases, enhancing the embryo’s directional response to stiffness cues. Critically, in our uterine scar model, we were able to significantly reduce implantation within scarred regions by inhibiting embryonic durotaxis using AACOCF3 pretreatment. These results establish that local uterine biomechanics directly regulate embryonic migration and implantation site selection, providing a mechanistic basis for the development of cesarean scar pregnancy (Supplementary Fig. S6).

Interestingly, collective migration behaviors have been described in tumor spheroids from lung, colorectal, and breast cancers (*48–51*). These malignant cell clusters often migrate cohesively through heterogeneous microenvironments, enhancing their metastasis. Thus, our findings reveal a fundamental mechanobiological mechanism that may extend beyond embryogenesis to broader contexts of collective cell migration in pathological states.

While our study sheds light on the role of stiffness gradients in guiding embryo migration and implantation, several limitations warrant consideration. First, our *in vitro* platform, comprising mouse embryos, human trophoblast spheroids, and stiffness-tunable polyacrylamide hydrogels, does not fully recapitulate the physiological complexity of the native uterine environment. To more accurately model trophectoderm–uterine interactions, future systems should incorporate additional components of the maternal interface, such as endometrial epithelial or stromal cells, within the hydrogel substrate. Second, although embryos in our system exhibited durotactic migration, the migration distance was still limited compared with the scale of embryo displacement observed in cesarean scar pregnancy. Previous studies have shown that uterine contractions and intraluminal fluid flow play important roles in regulating embryo transport in vivo (*13–15*), suggesting that such external forces may amplify durotaxis and enable longer-range migration in the uterine environment. Finally, although we used mouse embryos to investigate durotaxis, significant differences exist between murine and human implantation, particularly during later stages that involve trophoblast invasion and uterine tissue remodeling (*52–54*). Developing humanized in *vitro* models, either using human embryos or stem cell-derived trophoblast tissues, could help overcome this limitation and provide deeper mechanistic insight into human implantation and CSP pathogenesis. Such models would not only improve translational relevance but also enhance the potential for developing targeted therapeutic interventions for implantation-related disorders.

In summary, our findings establish that embryonic migration and adhesion are intrinsically mechanosensitive, governed by stiffness gradients within the microenvironment. These insights underscore the importance of biomechanical engineering approaches in reproductive biology, opening new directions for optimizing assisted reproductive technologies. More broadly, by uncovering mechanically regulated pathways in early development, our work lays the foundation for novel therapeutic strategies aimed at mitigating implantation-related complications. This integration of mechanobiology and reproductive medicine defines a new framework for bridging fundamental research with clinical applications in fertility and obstetrics.

## ACKNOWLEDGEMENTS

This work was supported by the National Key Research and Development Program of China (Nos. 2023YFF0613604, 2021YFA0719302, 2022YFF0503504), National Natural Science Foundation of China (Nos. 12222201, 82273500, 12332019, 12472273, and under Grant No.12372267), Beijing Nature Science Foundation (No.7232208), Clinical Cohort Construction Program of Peking University Third Hospital (No.2022015), Fundamental Research Funds for the Central Universities (No. ZG140S1971), and the Strategic Priority Research Program of Chinese Academy of Sciences (Grant No. XDB0620102).

## AUTHOR CONTRIBUTIONS

J.D., Y.B.F., Y.W., and X.Z. designed and supervised the experiments and modeling. J.L.C., H.Y.Y., J.W.L. performed the experiments. H.Y. Y., S.P.Y., H.Y.L., J.Y.W., D.S.G., J.E.Z., and J.Q.W. performed the uterine surgical scarring mouse model assays and AFM experiments. Z.X. developed the computational model. All the authors took part in the data analysis. J.L.C., J.D., H.Y.Y., X.Z., and Y.Z. interpreted the data and wrote the paper.

## MATERIALS AND METHODS

### Fabrication of polyacrylamide hydrogels with stiffness gradients

Polyacrylamide (PA) hydrogels with stiffness gradients were fabricated to mimic the mechanical properties of uterine tissue. A polymer solution was prepared by mixing 20% (v/v) acrylamide monomer (Bio-Rad) and 0.4% (v/v) N, N’-methylenebisacrylamide cross-linker (Bio-Rad) in 1× Dulbecco’s phosphate-buffered saline (PBS, Life Technologies, Gaithersburg, MD, USA) without Mg²⁺ and Ca²⁺. The solution was degassed for 5 minutes, after which 4 µL of 10% (w/v) ammonium persulfate (APS, Sigma-Aldrich) was added and mixed for 0.5 seconds. Subsequently, 1.2 µL of N, N, N’, N’-tetramethylethylenediamine (TEMED, Bio-Rad) was rapidly added and mixed for 1 second. A 120 µL aliquot of the solution was transferred into a glass mold, and a functionalized glass coverslip was placed over the solution at an angle to prevent air bubbles. Polymerization was allowed to proceed for 30 minutes at room temperature, after which the coverslip was removed. The gels were rinsed and stored in 1× PBS without Mg²⁺ and Ca²⁺ for later use.

To enable cell adhesion, the PA gel surface was functionalized using Sulfosuccinimidyl 6-(4’-azido-2’-nitrophenylamino) hexanoate (sulfo-SANPAH, Thermo Fisher Scientific, Waltham, MA, USA). The gel was exposed to sulfo-SANPAH for 15 minutes under 365 nm UV light (twice) and then coated with collagen I (100 µg/mL) overnight at 4 °C. Prior to cell seeding, the PA gels were washed three times with PBS. These PA hydrogels exhibited a thickness gradient and an apparent stiffness gradient, as previously described (*55*). The thickness of the PA gel was measured by staining the gel with Coomassie brilliant blue G-250 for 10 minutes. Z-stack images were acquired using a confocal microscope (Leica TCS SP8 X, Germany) and reconstructed to determine the gel’s thickness. Due to the minimal slope of the gel (< 1⁰), the gravitational effect is negligible.

The surface topography of the PA gel was examined using scanning electron microscopy (SEM, Quanta 200 FEG). The gels were flash-frozen and lyophilized overnight, followed by sputter-coating with a layer of platinum (Pt) using a turbomolecular pump coater (Q150T, Quorum Technologies, Lewis, UK) to enhance electrical conductivity. Each PA gel was fabricated under consistent conditions across independent experiments, and reproducibility was verified by controlling the size, thickness, and partitioning of the gels.

### Mice

All studies involving animals have been reviewed and approved by the Peking University Third Clinical Medical School Ethical Committee of animals (Approval No. A2023001). Eight-week-old CD-1 (CD-1 IGS) mice were purchased from Charles River (Beijing, China). Both male and female mice were purchased and housed in a specific pathogen-free (SPF) animal facility under standard environmental conditions, with constant temperature, humidity and 12h: 12h light cycle. The food and water were available ad libitum.

In this study, pregnancies were created using timed mating. One or two females in estrus were placed with a stud male overnight. Visualization of copulation plug was denoted as embryonic day 0.5 (E0.5), Successfully mated females were weighted and removed from the stud cage.

### Embryo collection and culture

Preimplantation embryos were collected from female mice (aged 2 to 3 months) to study their behavior in vitro. To induce superovulation, female mice were intraperitoneally injected with 10 international units (IU) of pregnant mare serum gonadotropin (PMSG; Solarbio, P9970). Forty-eight hours later, the mice were injected with 10 IU of human chorionic gonadotropin (hCG; ShuSheng, 11025). Immediately after the hCG injection, the female mice in the superovulation phase were mated with male mice (aged 3 to 4 months). Preimplantation embryos were collected by flushing the uterus or fallopian tubes with KSOM medium (Caisson, IVL04). The collected embryos were then cultured in drops of KSOM under mineral oil (Sigma, M8410) at 37 °C in a 5% CO₂ atmosphere. Both the mineral oil and KSOM medium were preheated in the incubation chamber for at least 30 minutes prior to embryo culture to ensure optimal conditions. All embryo manipulations were performed using a stereo microscope (Olympus SZX16).

At E3.5, the zona pellucida was removed using acidic Tyrode’s solution (Sigma, T1788), and embryos were transferred into M2 medium (Sigma, M7167) before returning to KSOM. Embryos were then cultured until E4.5 and subsequently placed onto stiffness-gradient hydrogels for in vitro implantation experiments. Embryos were maintained in IVC1 medium (Advanced DMEM/F-12 supplemented with 20% FBS, 2 mM L-glutamine, 25 U/mL penicillin and 25 µg/mL streptomycin, 1× ITS-X, 8 nM β-estradiol, 200 ng/mL progesterone, and 25 µM N-acetyl-L-cysteine) at 37 °C in 5% CO₂.

### General cell culture

The JAR human choriocarcinoma cell line (TCHu156, ATCC) was maintained in RPMI 1640 medium (Hyclone) at 37°C in a humidified incubator with 5% CO₂. The culture medium was supplemented with 10% (v/v) fetal bovine serum (FBS, SERANA), 100 units/mL penicillin, and 100 units/mL streptomycin.

### JAR spheroid formation

For the spheroid formation assay, JAR cells were seeded at a density of 1 × 10^5^ cells/mL in 100 µL per well in an ultra-low attachment 96-well round-bottom plate (Corning, NY, USA). Cells were incubated overnight at 37 °C in a 5% CO₂ incubator. During this incubation, JAR cells naturally aggregated to form spheroids with diameters ranging from 50 to 200 µm. The aggregate diameters of the spheroids were measured using CellSens standard software (Olympus, Tokyo, Japan).

### Uterine scar mouse model and uterine perfusion experiments

Virgin mice were used. Mice were anesthetized with isoflurane inhalation anesthesia using a precision vaporizer. 2.5% isoflurane was used for induction, then reduced to 1.5% for maintenance anesthesia throughout the surgery via nosecone. And the oxygen flow was set at 1.5 L/min. A low abdominal incision was made to expose the uterine horns. Chose the right or left uterine horn randomly, and a longitudinal incision of the anti-mesometrial side was made from the cervical side of the uterine horn to the midpoint of the uterine horn. The incision length was half the length of the chosen uterine horn. The incision penetrated through the myometrium and endometrium of the anti-mesometrial side, but did not harm the mesometrial side. The incision was closed with 8-0 absorbable sutures using running locked suture. The end near the midpoint of the uterine horn was marked with 4-0 nonabsorbable sutures. The uterine was placed back into the abdominal cavity, and the abdominal wall and skin were closed by simple continuous suture with 5-0 absorbable suture respectively. Following surgery, the mice were permitted to recover for 20 days and mated to fertile male mice. The pregnant mice were humanely killed at different embryonic day.

For mouse uterine perfusion experiments, pregnant mice at embryonic day 3.5 (E3.5) were anesthetized via intraperitoneal injection of tribromoethanol (Avertin, 270–330 mg/kg). Mice were positioned supine with limbs taped to the operating platform, which was tilted 60–70° to elevate the caudal end and facilitate cervical access. A 1 mL pipette tip was used to fabricate a vaginal dilator, which was inserted into the mouse’s vagina and gently advanced to reach the cervical os with extreme care. A disposable side-opening dental irrigation needle was advanced into one uterine horn for unilateral perfusion. Once intubation was completed, one side of the uterus was subjected to perfusion for drug administration, with a perfusion volume of 10-15 μL. During perfusion, attention was paid to the flow rate to prevent fluid leakage. The same perfusion procedure was followed for the other side of the uterus. After perfusion, the mice were maintained in an inverted position for 1-2 minutes to prevent the perfusate from flowing out. Meanwhile, dorsal and abdominal massage was performed to achieve uniform distribution of the perfusate.

### Human tissues

Samples of human uterine were obtained from patients who had undergone hysterectomy at the Obstetrics and Gynecology Department of Peking University Third Hospital. The patients had a prior history of cesarean section. The obtained samples consisted of partial anterior wall tissue of the uterus. This tissue collection did not affect the patient’s diagnostic evaluation or therapeutic care, as it represented clinical residual biospecimen. All patients provided written informed consent. This study received approval from the Ethics Committee of Peking University Third Hospital (IRB00006761-M2023275).

### Hematoxylin-eosin and immunohistochemistry staining

The tissues were fixed with 4% paraformaldehyde and embedded in paraffin. Tissue sections (4 μm thick) were stained with hematoxylin and eosin (HE). For immunohistochemistry (IHC) staining, after antigen repair, the tissue sections were incubated in darkness at room temperature in 3 % H_2_O_2_ solution for 15 min. Then, the sections were coated with the following solution of primary antibodies and incubated at 4 °C overnight: α-SMA (1:800, AFMM0002, Aifang Biotechnology). Subsequently incubated with a solution of horseradish peroxide-coated secondary antibody at 37 °C for 50 min, then developed with DAB substrate (AFIHC004, Aifang Biotechnology) to visualize the antigen signals and counterstained with hematoxylin. All sections were examined under an E100 Nikon microscope. For IHC result analysis, relevant regions were traced and measured using K-Viewer software. Semi-automated quantification of positive immunoreactivity intensity was performed using ImageJ 1.54g.

### Embryo labeling and imaging

For nuclear labeling, embryos were incubated with Hoechst 33342 (Thermo Fisher, H1399) at 5 µg/mL for 5–10 min in IVC1 medium followed by washes, or with SPY555-DNA (Cytoskeleton, SC201) at 1:1000 dilution in IVC1 medium for ∼1 h at 37 °C and 5% CO₂ prior to imaging. For real-time imaging, embryos were cultured in IVC1 medium on stiffness-gradient hydrogels in 35 mm glass-bottom dishes and maintained at 37 °C in 5% CO₂. Time-lapse imaging was performed using either an Andor Dragonfly Fusion 2.2 spinning-disk confocal microscope (Oxford Instruments) or a Cytation C10 automated confocal imaging system (BioTek), both equipped with temperature- and CO₂-controlled environmental chambers.

### Atomic Force Microscope (AFM) measurement

The apparent Young’s modulus of the PA gel was assessed using an atomic force microscope (AFM, MFP-3D, Asylum Research Inc., Santa Barbara, CA, USA; AFM AtomEdge Pro from Truth Instruments Company (Qingdao) equipped with a colloidal probe cantilever. A non-tip cantilever (NSC36, MikroMasch, Watsonville, CA, USA) with a spring constant of 2 N/m was used. The colloidal probe, consisting of a glass sphere (d = 26.3 µm) adhered to the cantilever’s front end, was prepared as described previously (*56*). To minimize adhesion between the probe and the PA gel, the colloidal probe surface was coated with PLL-g-PEG (SuSoS AG) (*57*). AFM measurements were performed in contact mode with indentation depths of 1–8 µm. Force-distance curves were recorded and analyzed using the Hertz model 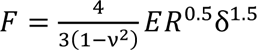 to determine the reduced Young’s modulus *E* (*58*), where *F* is the force, *R* is the probe radius, δ is the indentation distance, and ν = 0.33 represents the Poisson ratio. The spring constant of the colloidal probe was calibrated in situ before each measurement using the thermal power spectral density method (*56*).

For uterine tissue measurements, pregnant female mice with scarred uteri were sacrificed at E3.5. The uteri were dissected, longitudinally opened along the antimesometrial side, and fixed flat on glass-bottom dishes. AFM indentation experiments were performed on the exposed endometrial surface using identical parameters and analysis procedures as for PA hydrogels.

### FLIM acquisition and analysis

For Flipper-TR staining, mouse embryos were incubated with 1 μM Flipper-TR (SC020, Spirochrome) at 37 °C in a humidified atmosphere containing 5% CO_2_ for 15 minutes and transferred to PA gel in 35mm glass-bottom petri dishes before imaging. Imaging was performed using Olympus fluorescence lifetime imaging microscope (FLIM, FV-1200, Japan). Excitation can be commonly performed with a 488nm laser, while emission is collected between 575 and 625nm bandpass filter using a HyD SMD detector. SymPhoTime 64 software (PicoQuant) was used to analyze FLIM data. The fluorescence decay data were fitted with a double-exponential. Two decay times were extracted as τ1 and τ2. The longest lifetime with the higher fit amplitude τ1 was used to report membrane tension.

### Particle Image Velocity (PIV) measurement

Particle Image Velocimetry (PIV) analysis was conducted using a custom algorithm based on MATLAB’s PIVlab software package (Matlab2020a, PIVlab2.36). Live cell image sequences of JAR cells were used to analyze the direction and magnitude of cell movement. To mitigate the influence of background movement on the calculated results, the average background velocity was subtracted from the calculated velocity field. For the velocity vector arrows, each pixel length corresponds to 0.05 µm/min.

### Statistical analysis

All statistical analyses were conducted using R software (version 4.1.1). Data are presented as mean ± standard deviation (SD). Each experiment was independently repeated three times (n = 3). Prior to statistical testing, data normality and homogeneity of variances were assessed using the Shapiro–Wilk test and Bartlett’s test, respectively. For comparisons between two groups, Student’s t-test was applied. A p-value less than 0.05 was considered statistically significant.

## SUPPLEMENTAL FIGURES

**Fig. S1.**
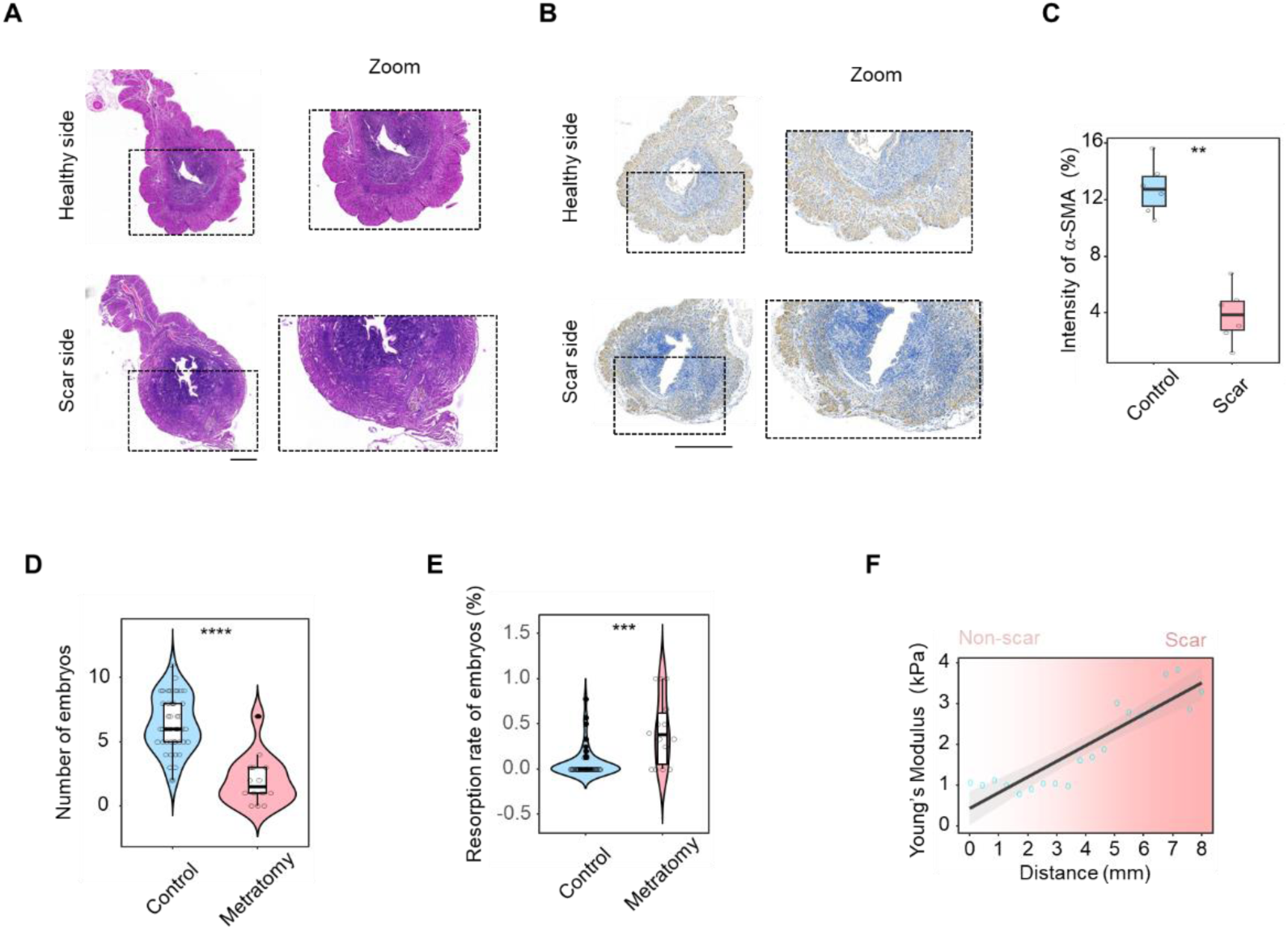
Histological analysis of murine uterine scar model. (**A**) Representative images of HE staining in murine uterine scar model. Scale bar: 500 µm. (**B**) Representative images of immunohistology analysis of α-SMA in a murine uterine scar model. Scale bar: 800 µm. (**C**) Quantification of α-SMA intensity in the myometrium of the antimesometrial side or scar area. (**D**) The statistical analysis of embryo number at E7.5 stage in mice with metratomy or control mice, n = 44 (control), 14 (metratomy). (**E**) The statistical analysis of embryo resorption rate in mice with metratomy or control mice, n = 44 (control), 14 (metratomy). (**F**) The AFM analysis of Young’s modulus from non-scarred to scarred regions in the scarred human uterine. * p < 0.05, ** p < 0.01, *** p < 0.001, **** p < 0.0001.

**Fig. S2.**
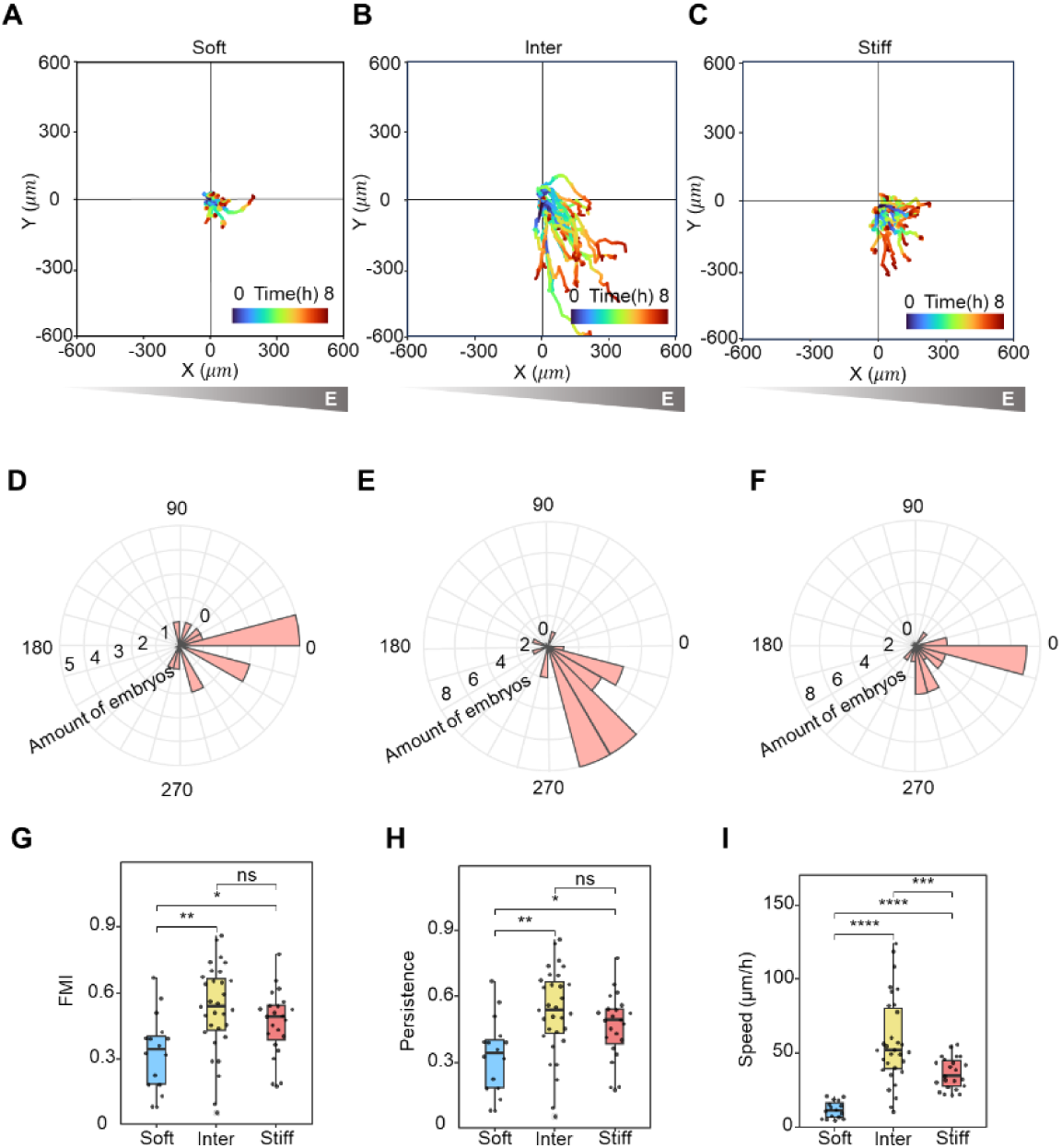
Embryonic durotaxis on substrates with gradient stiffness at different stiffness zones. (**A-C**) Embryo migration plots on gradient substrates over 8 h on substrates with gradient stiffness at different stiffness zones. (**D-F**) Rose diagram of embryo migration direction on substrates with gradient stiffness at different stiffness zones, which displays the angular between migration and stiffness gradient and the frequency of each class. (**G-I**) The statistical analysis of durotaxis-related parameters, including Forward Migration Index (FMI), persistence, and velocity of embryonic migration on substrates with gradient stiffness at different stiffness zones. All box-and-whisker plots show the medians, maxima, minima, upper quartiles, and lower quartiles, n = 16 (soft), 31 (inter), 23 (stiff). * p < 0.05, ** p < 0.01, *** p < 0.001, **** p < 0.0001.

**Fig. S3.**
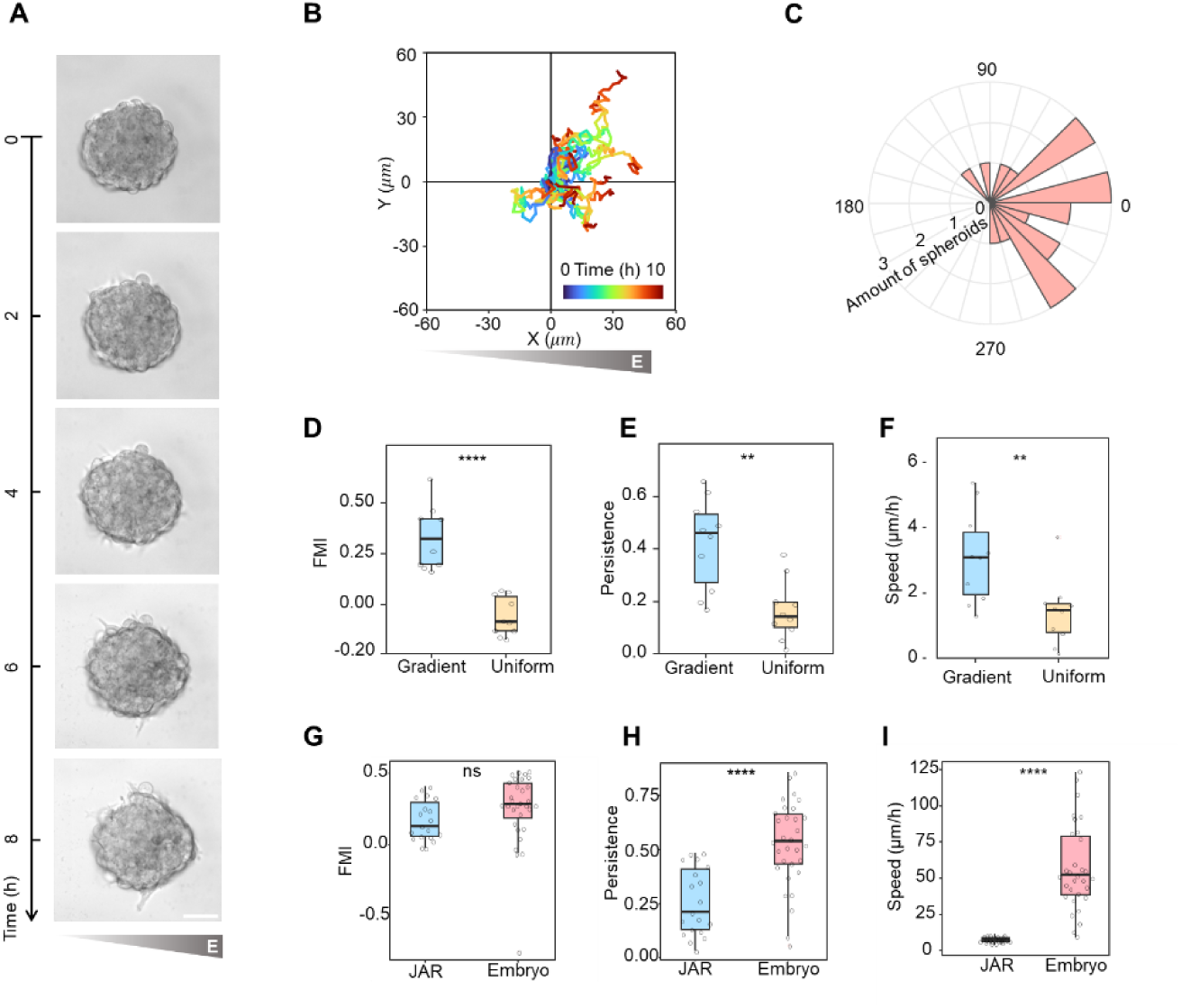
Durotaxis of human trophoblast spheroids. (**A**) Representative time-lapse images of JAR spheroid on substrates with uniform stiffness or gradient stiffness. Scale bar: 20 μm. (**B**) JAR spheroid migration plots on gradient substrates over 10 h. (**C**) Rose diagram of JAR spheroid migration direction on substrates with uniform stiffness or gradient stiffness, which displays the angular between migration and stiffness gradient and the frequency of each class. (**D-F**) The statistical analysis of durotaxis-related parameters, including Forward Migration Index (FMI), persistence, and velocity of JAR spheroid migration on substrates with uniform stiffness or gradient stiffness, n = 9. (**G-I**) The statistical analysis of durotaxis-related parameters, including Forward Migration Index (FMI), persistence, and velocity of JAR spheroid or embryo migration on substrates with uniform stiffness or gradient stiffness, n = 19. All box-and-whisker plots show the medians, maxima, minima, upper quartiles, and lower quartiles. * p < 0.05, ** p < 0.01, *** p < 0.001, **** p < 0.0001.

**Fig. S4.**
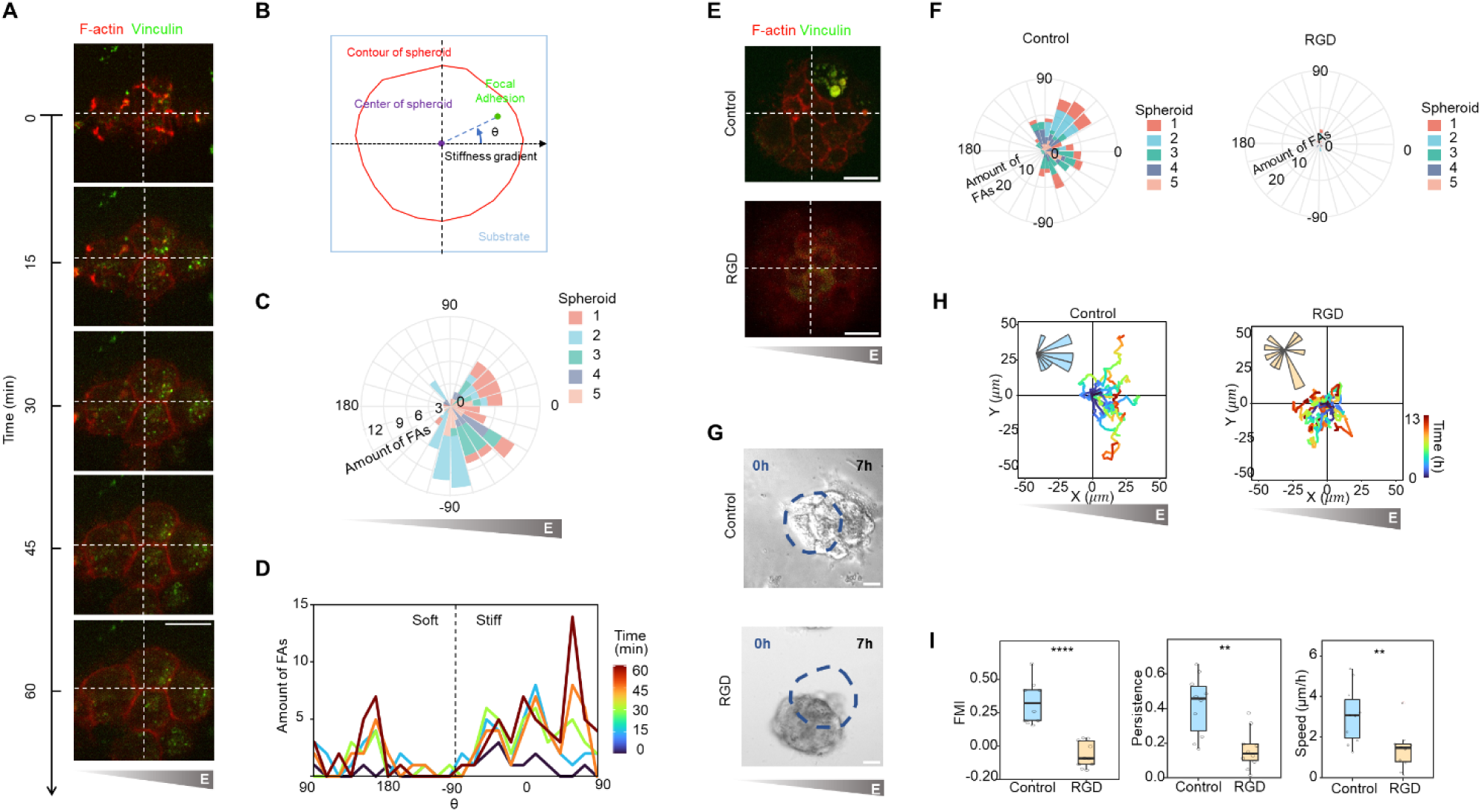
The establishment of an asymmetric cell-substrate adhesion pattern is required for durotaxis of human trophoblast spheroids. (**A**) Representative time-lapse fluorescent images of Vinculin-GFP and Lifeact-mCherry signals at the spheroid-substrate interface in Vinculin-GFP^+^/Lifeact-mCherry^+^ JAR spheroids. Scale bar: 20 μm. (**B**) Schematic illustration of angles (θ) between membrane signal related to spheroid center and substrate stiffness gradient. (**C**) Rose diagram of Vinculin-GFP signals at the spheroid-substrate interface in Vinculin-GFP^+^/Lifeact-mCherry^+^ JAR spheroids, which displays angles (θ) between Vinculin-GFP signal related to spheroid center and substrate stiffness gradient. (**D**) The Vinculin-GFP intensity at the spheroid-substrate interface in Vinculin-GFP^+^/Lifeact-mCherry^+^ JAR spheroids at different angles (θ) during the adhesion of spheroids to substrates. (**E**) Representative time-lapse fluorescent images of Vinculin-GFP and Lifeact-mCherry signals at the spheroid-substrate interface in Vinculin-GFP^+^/Lifeact-mCherry^+^ JAR spheroids treated with control or RGD. Scale bar: 20 μm. (**F**) Rose diagram of Vinculin-GFP signals at the spheroid-substrate interface in Vinculin-GFP^+^/Lifeact-mCherry^+^ JAR spheroids treated with control or RGD, which displays angles (θ) between Vinculin-GFP signal related to spheroid center and substrate stiffness gradient. (**G**) Representative time-lapse images of JAR spheroids on stiffness gradient substrates treated with control or RGD. Scale bar: 20 μm. (**H**) JAR spheroid migration plots on stiffness gradient substrates over 7 h treated with control or RGD. Insets are rose diagram of spheroid migration direction, which displays the angular between migration and stiffness gradient and the frequency of each class. (**I**) The statistical analysis of durotaxis-related parameters, including Forward Migration Index (FMI), persistence, and velocity of JAR spheroid migration on stiffness gradient substrates over 7 h treated with control or RGD, n = 10 (control), 9 (RGD). All box-and-whisker plots show the medians, maxima, minima, upper quartiles, and lower quartiles. * p < 0.05, ** p < 0.01, *** p < 0.001, **** p < 0.0001.

**Fig. S5.**
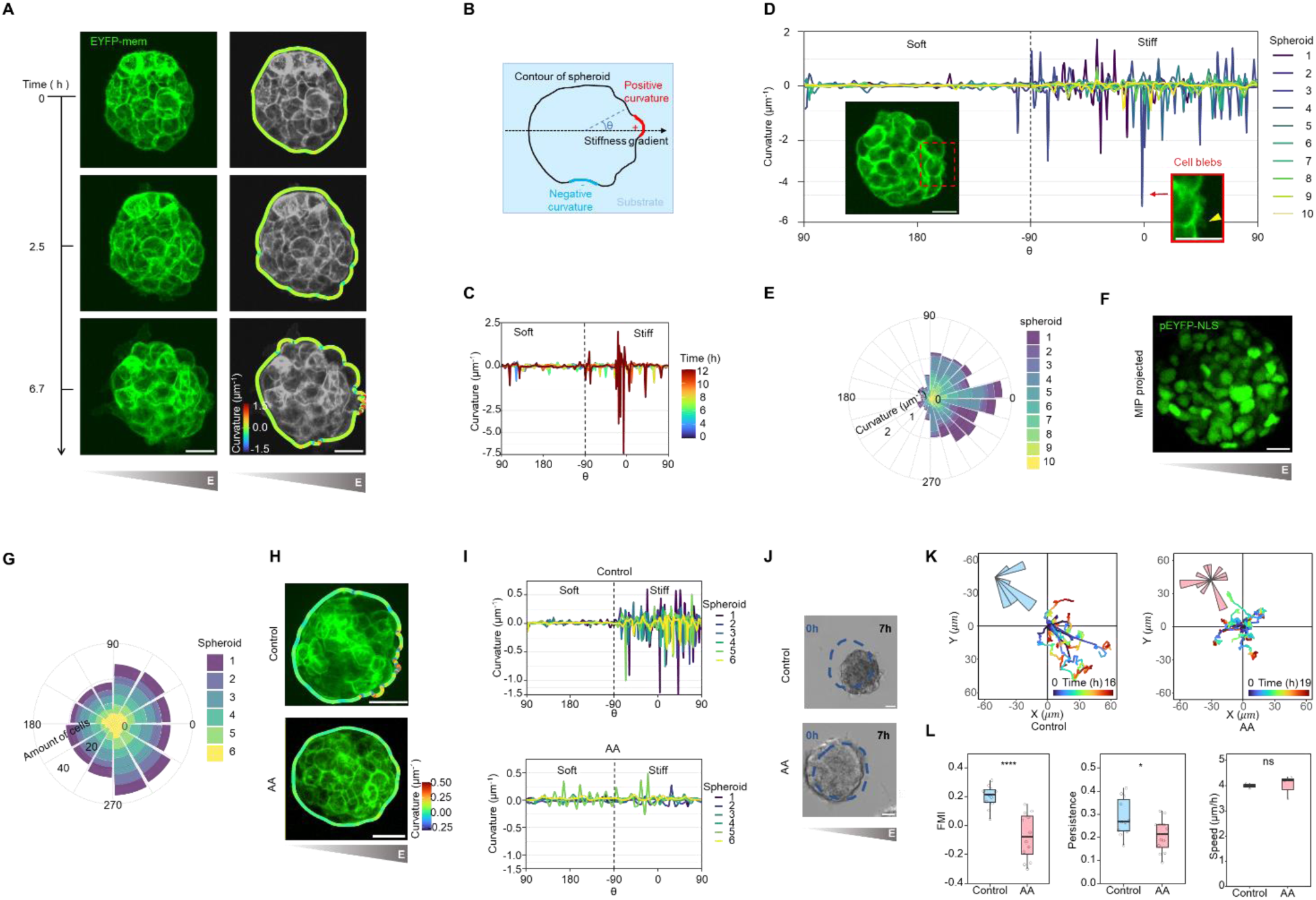
Inhibiting cell blebbing disrupts durotaxis of human trophoblast spheroids. (**A**) Representative time-lapse images of surface morphology in EYFP-mem^+^ JAR spheroids over time on substrates with gradient stiffness. Scale bar: 20 μm. (**B**) Schematic illustration of angles (θ) between surface curvature to spheroid center and substrate stiffness gradient. (**C**) The surface curvature of JAR spheroids at different angles (θ) over time on substrates with gradient stiffness. (**D**) The surface curvature of JAR spheroids at different angles (θ) on substrates with uniform stiffness or gradient stiffness. Insets are presentative fluorescent images of EYFP-mem^+^ JAR spheroids with two distinct types of surface fluctuations. Scale bar: 20 μm. (**E**) Rose diagram of surface curvature of JAR spheroids, which displays angles (θ) between membrane signal related to spheroid center and substrate stiffness gradient. (**F**) Representative fluorescent images of cell nuclei in EGFP-NLS^+^ JAR spheroids during durotaxis on stiffness gradient substrates. Scale bar: 20 μm. (**G**) The cell density at different angles (θ) on substrates with gradient stiffness. (**H**) The analysis of JAR spheroid surface curvature on stiffness gradient substrates treated with control or AACOCF3. Scale bar: 20 μm. (**I**) The surface curvature of JAR spheroids at different angles (θ) on stiffness gradient substrates treated with control or AACOCF3. (**J**) Representative time-lapse images of JAR spheroids on stiffness gradient substrates treated with control or AACOCF3. Scale bar: 20 μm. (**K**) JAR spheroid migration plots on stiffness gradient substrates treated with control or AACOCF3. Insets are rose diagram of spheroid migration direction, which displays the angular between migration and stiffness gradient and the frequency of each class. (**L**) The statistical analysis of durotaxis-related parameters, including Forward Migration Index (FMI), persistence, and velocity of JAR spheroid migration on stiffness gradient substrates treated with control or AACOCF3, n = 11 (control), 12 (AA). All box-and-whisker plots show the medians, maxima, minima, upper quartiles, and lower quartiles. * p < 0.05, ** p < 0.01, *** p < 0.001, **** p < 0.0001.

**Fig. S6.**
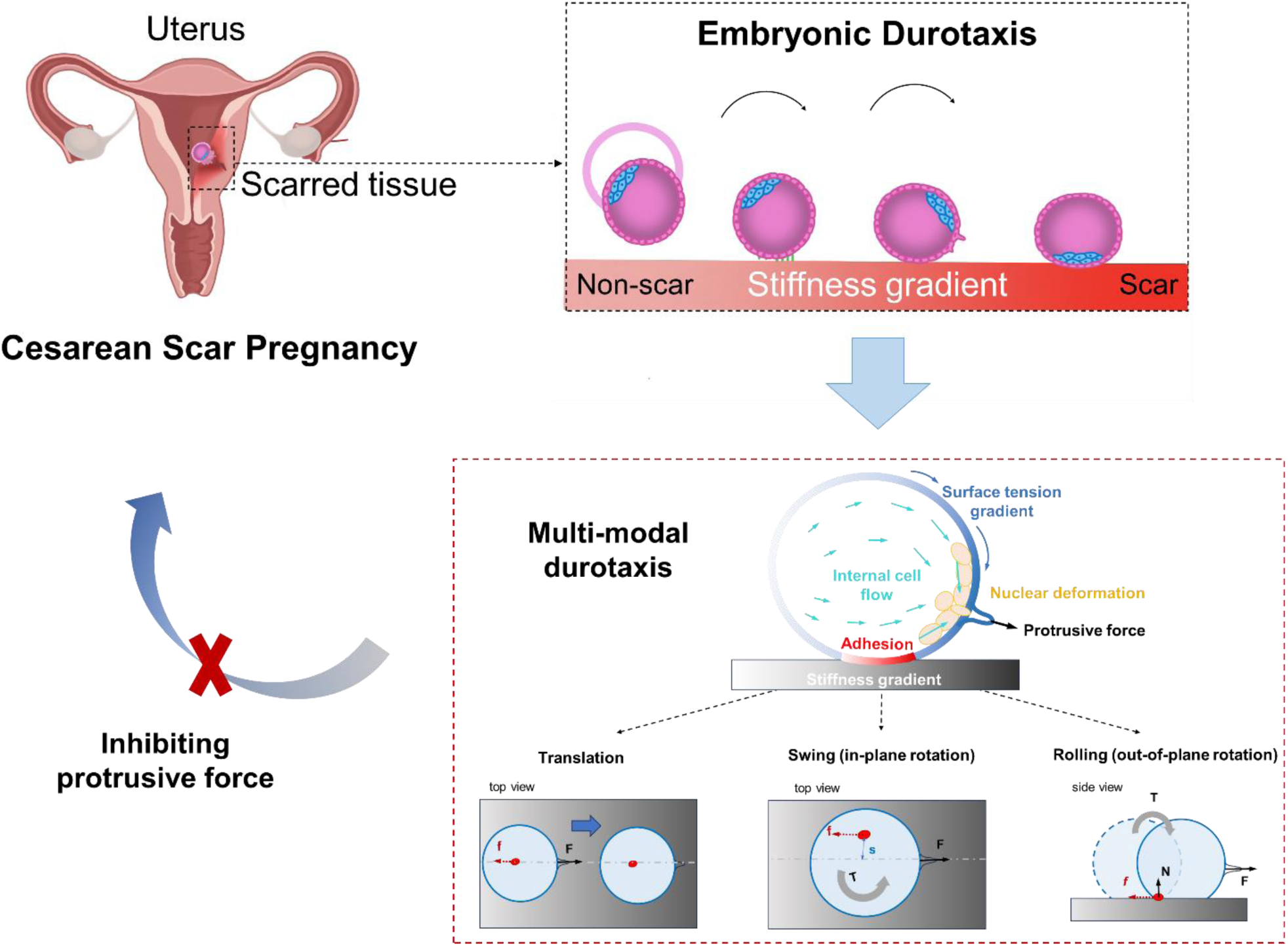
The model of mammalian embryonic durotaxis and its role in CSP pathogenesis. Mammalian embryos undergo durotaxis, migrating directionally along uterine stiffness gradients from non-scarred to scarred regions post-cesarean section. This process is driven by polarized trophectoderm cell blebbing, generated by internal tissue flows toward stiffer side and manifests in three kinematic modes: translational, swing, and rolling. Disrupting this blebbing-mediated force inhibits durotaxis, reducing implantation in scarred regions and highlighting its role in CSP pathogenesis.

